# The role of *AUX1* during lateral root development in the domestication of the model C4 grass *Setaria italica*

**DOI:** 10.1101/2021.05.20.444970

**Authors:** Sha Tang, Mojgan Shahriari, Jishan Xiang, Taras Pasternak, Anna Igolkina, Somayeh Aminizade, Hui Zhi, Yuanzhu Gao, Farshad Roodbarkelari, Yi Sui, Guanqing Jia, Chuanyin Wu, Xugang Li, Georgy Meshcheryakov, Maria Samsonova, Xianmin Diao, Klaus Palme, William Teale

## Abstract

C4 photosynthesis increases the efficiency of carbon fixation by spatially separating high concentrations of molecular oxygen from rubisco. The specialized leaf anatomy required for this separation evolved independently many times. C4 root systems are highly branched, an adaptation thought to support high rates of photosynthesis; however, little is known about the molecular mechanisms that have driven the evolution of C4 root system architecture (RSA). Using a mutant screen in the C4 model plant *Setaria italica*, we identify *siaux1-1* and *siaux1-2* as RSA mutants, and use CRISPR/cas9-mediated genome editing and overexpression to confirm the importance of the locus. As *AUX1* is not necessary for lateral root emergence in *S. viridis*, the species from which *S. italica* was domesticated, we conducted an analysis of auxin responsive elements in the promoters of auxin-responsive gene families in *S. italica*, and explore the molecular basis of *SiAUX1’*s role in seedling development using an RNAseq analysis of wild type and *siaux1-1* plants. Finally, we use a root coordinate system to compare cell-by-cell meristem structures in *siaux1-1* and wild type *Setaria* plants, observing changes in the distribution of cell volumes in all cell layers and a dependence in the frequency of protophloem and protoxylem strands on *siAUX1*.

## Introduction

Improving and adapting plant growth is central to many of the world’s most pressing challenges. For example, there is a broad consensus that the world’s hydrocarbon economy must switch rapidly from the release of carbon from oil and gas reserves to the capture and release of energy through the cycling of existing atmospheric CO_2_ (Koçar and Civas, 2013). However, not all plants sequester CO_2_ with equal efficiency; photorespiration, a process that is initiated by the reaction of ribulose bisphosphate with O_2_ instead of CO_2_, acts as a major and highly variable limiting factor (Sage et al., 2012). In response, many of the species which carry the most potential for use in the production of bioenergy such as switchgrass and *Miscanthus* employ C4 photosynthesis, maintaining relatively high CO_2_:O_2_ ratios in CO_2_-fixing cells (Kocar and Civas, 2013).

Fast growth rates in C4 species place a high demand on root systems for resources. Key features of root systems that have evolved in response to fast C4 growth have been identified after a careful comparative analysis of growth patterns in different grass species (Wade et al., 2020). An important common feature of these C4 grasses is an increased rate of root branching when compared to C3 grasses. However, it is not possible to infer a common molecular C4 module that has driven increased branching rates, as C4 photosynthesis evolved at least 66 times independently in the last 35 million years (Sage et al., 2012). It is likely, though, that auxin will be an important factor in driving root system architecture as it is a key regulator of root system development throughout the embryophyte clade (Yu et al., 2016; Péret et al., 2009).

In many key developmental contexts, the cellular concentration of auxin is largely determined by regulated transport via influx and efflux carriers (Teale et al., 2006). One such protein, AUXIN PERMEASE 1 (AUX1) is an auxin influx carrier that is particularly relevant to root development (Perét et al., 2009). In Arabidopsis, loss of *aux1* leads to a distinctive range of phenotypes, including agravitroptic growth and the inability to grow lateral roots, (Bennett et al, 1996).

An important model C4 genus is *Setaria*: either the wild species *S. viridis* or its domestic relative *S. italica*. Its use has been driven by important, foundation-laying work in, for example, the development of transformation protocols (Brutnell et al, 2010), and the availability of reference genomes (Bennetzen et al, 2012; Zhang et al, 2012). *Sparse panicle1* (*spp1*) is an AUX1 loss of function allele in *Setaria viridis* which was identified as a mutant that is severely affected in the development of its inflorescence (Huang et al., 2017a). However, although *spp1* displays the same agravitropic root phenotype observed in Arabidopsis, it is unusual among *aux1* loss-of-function mutants as is not deficient in lateral root development (Huang et al., 2017a).

In this report, we generate loss-of-function *aux1* alleles of *S.italica* to examine the role of SiAUX1 in shaping roots system architecture throughout domestication. Although *S. viridis spp1* plants remain unaffected in their ability to produce lateral roots, *Siaux1-1 S. italica* plants produces lateral roots at a reduced frequency, indicating that *AUX1*-independent lateral root suppression may have been overwritten during domestication. We used RNAseq to explore the molecular basis for this difference, suggesting a small subset of genes that could be involved in the SiAUX1-dependent determination of root development. Using a high-resolution phenotyping platform to perform a cell-by-cell analysis of the *Setaria italica* root apical meristem, we also examine the role SiAUX1 plays in shaping the root apical meristem, finding it affects cell volumes and vascular structure.

## Materials and methods

### Mutant isolation and phenotype characterization

The *Setaria italica* sparse panicle mutants *siaux1-1* and *siaux1-2* were screened out from an EMS-induced mutant library based on a modern variety, Yugu1, with an available, full genome sequence (https://phytozome.jgi.doe.gov/pz/portal.html#!info?alias=Org_Sitalica). For the phenotyping of major agronomic traits, mutants and wild-type plants were grown in experimental plots at the Institute of Crop Science, Chinese Academy of Agricultural Sciences, during the foxtail millet growth period from June to October in 2017 (Beijing, China). Measurements of agronomic traits including plant height, panicle length, leaf length, were done according to our previous study (Jia et al, 2013). For scanning electron microscope (SEM) analysis, young panicles (less than 0.5 cm long) of *siaux1-1* and Yugu1 were fixed and dehydrated according to a previous report (Hodge & Kellogg, 2014). Observation was conducted using a Hitachi S3400N SEM (Japan). For characterization of root development, *siaux1-1* and Yugu1 seeds were germinated on damp flat filter paper, and then cultivated in water for 10 days before analysis. Phosphate-depleted medium contained 10mM NH_4_NO_3_, 9.5 mM KNO_3_, 1mM CaCl_2_, 720 μM MgSO_4_, 50 μM BH_3_O_3_, 40 μM ZnNa_2_-EDTA, 25 μM FeCl_3_, 15 μM ZnSO_4_, 2.5 μM KI, 0.75 μM MoO_3_ 50 nM CoCl_2_, 50 nM CuSO_4_; 1-naphtheleneacetic acid (1-NAA) was applied in solid medium at 100 nM.

### Map-based cloning of SiAUX1

The ‘*siaux1-1’* mutant was crossed with ‘SSR41’ (a foxtail millet variety which shared high polymorphism with ‘Yugu1’) to obtain the F_2_ mapping population. About 540 homozygote recessive F_2_ plants were collected and used for DNA extraction. PCR-based mapping was conducted using the molecular markers listed in Supplementary Table S3.

### Whole genome resequencing and mutmap analysis for S.italica aux1 mutants

The ‘*siaux1-1’* mutant and the ‘*siaux1-2’* mutant were both backcrossed with their parental line ‘Yugu1’ to obtain the BC_1_F_2_ population. Genetic analysis was performed through the analysis of segregation ratios. The result (Supplementary Table S1) indicated that the *Setaria aux1* phenotype was recessive. We then collected leaf samples from individual recessive plants from ‘*siaux1-1*’×‘Yugu1’ and ‘*siaux1-2*’×‘Yugu1’ BC_1_F_2_ populations. 30 individuals from each population were used to make DNA pools, which were then submitted for whole-genome resequencing using an Illumina Hiseq 2500 in PE150 sequence mode. Clean reads were uploaded to EBI with accession number PRJEB35973. Mutmap analyses were done according to a previous report (Abe et al, 2012).

### F_1_ complementation testing

Analysis of complemented line was used to test whether *siaux1-1* and *siaux1-2* represented alleles of *SiAUX1*. The *siaux1-1* mutant was crossed with *siaux1-2* and F1 plants were harvested. Mutation sites in both *siaux1-1* and *siaux1-2* were amplified and sequenced in all of *aux1*-like F_1_ plants to identify the F_1_ hybrids.

### Plasmid Construction and Generation of Transgenic Plants

To generate a *pUbi:AUX1* overexpression plant, a CDS fragment of the *SiAUX1* gene (Seita.5G387200) was amplified from cDNA synthesized from wild-type Yugu1 using the primers (OE-AUX1) listed in Supplementary Table S3. A modified pCAMBIA-1300 vector was used to make the *pUbi:AUX1*-containing plasmid. *SiAUX1* gene knockouts were constructed in foxtail millet usingthe pYLCRISPR/Cas9-MH vector as previously described (Ma et al, 2015). Primers of CRIPSR vector construction and sgRNA sequence are shown in Supplementary Table S3.

All transgenic vectors were transferred to the *Agrobacterium* EHA105 strain, and calli of the foxtail millet Ci846 (A high transgenic efficiency genotype) were used for *Agrobacterium*-mediated transformation (Chen QN, 2018)(48). All transgenic plants were grown in a growth chamber under short day conditions (31℃/25℃ Day 10 hours/night 14 hours 40% humidity).

### Tissue sampling and RNAseq analysis

S*iaux1-1* and wild type Yugu1 plants were grown in a soil mixture containing packaged potting soil, peat moss and sand (2:2:1, v/v/v) in a growth chamber (10 hours with 30°C in Day to 14 hours with 26°C as night). In total, 10 tissue samples were collected for RNA sequencing in each genotype. These samples included leaf and root tissues from seedling plants with either one fully expanded leaf (1 leaf stage), three fully expanded leaves (3 leaf stage), a young panicle, or at heading. All tissues were immediately frozen in liquid nitrogen and stored at −80°C. RNA extraction was preformed using Trizol (Ambion, Thermo Scientific Inc., USA) with three independent replications for each tissue. RNA samples were sent to the Beijing Berry Genomic facility (Beijing, China) for transcriptome sequencing. Thirty cDNA libraries were constructed and sequenced on an Illumina HiSeq X ten platform in 150-bp paired-end mode (Illumina Inc., USA). Sequencing data obtained in this study have been deposited at EMBL-EBI (http://www.ebi.ac.uk/) with the accession number PRJEB44706. The foxtail millet genome Yugu1 version 2.2 was used as reference genome (https://phytozome.jgi.doe.gov/pz/portal.html#!info?alias=Org_Sitalica). The RNA-seq data was analyzed as described in our previously (Tang et al., 2017).

### Searching for homologs in seven gene families

To reveal the correspondence between *Arabidopsis* and *Setaria* among genes of auxin pathway, we considered all members in seven target gene families: *AUX/LAX*, *GH3*, *AUX/IAA*, *PIN*, *ARF*, *LBD*, and *SAUR*. First, we searched the scientific literature for members of these families in Arabidopsis (Chettoor & Evans, 2015; Kakei et al, 2015; Kong et al, 2017; Markakis et al, 2013; Okrent & Wildermuth, 2011; Peret et al, 2012; Remington et al, 2004) and obtained the respective predicted protein sequences from the *Arabidopsis* genome (Krishnakumar et al, 2015): the *AUX/LAX* family contains 4 homologs, *ARF* – 23, *PIN* – 8, *LBD* – 43, *SAUR* – 79, *AUX/IAA* – 29, and *GH3* – 19. For each gene family, members were aligned with MUSCLE (Edgar, 2004) and a profile-HMM on the obtained alignment was constructed using HMMER (Eddy, 2011).

To search homologs of the target families, we collected all protein sequences of five grass species from Phytozome (Goodstein et al, 2012): *Setaria italica* (SI), *Zea mays* (ZM), *Oryza sativa* (OS), *Sorghum bicolor* (SB), and *Brachypodium distachyon* (BD). Then, we applied the HMMER search tool employing obtained profile-HMMs to collected protein sequences. Sequences with E-values less than 10^-6^ were classified as homologs and used for further analysis.

To reconstruct the evolution of each gene family, we aligned all its homologs in six target species with MUSCLE and applied the Neighbor-joining algorithm in pairwise deletion mode and with the bootstrap support (1000 iterations). As an outgroup for each gene family, we utilized one homolog from *Selaginella moellendorffii* (SM).

### Searching for auxin response motifs

For each of the homologs found in Arabidopsis and *Setaria* (Supplementary Table S4), we scanned its 1kbp upstream region to find auxin response motifs. We considered two types of motifs – TGTCNN and TGTSTSBC – in both direct and reverse-complement orientations (Mironova et al, 2014). For each gene, we counted single motifs and their pairs spaced by between 0 and 9 nucleotides.

### High-resolution phenotyping analysis

*Setaria* seeds were washed for three minutes in 70% ethanol, before sterilizing them for 10 minutes in a sonicating waterbath in 10% household bleach (aq), before washing ten times in sterile water and planting on ½ MS plates (1.3% agar). Three-day old seedlings were fixed in MTSB buffer containing 4% Formaldehyde and 0,3% Triton X100 for 45 minutes, a vacuum was applied after three minutes.

Seedlings were washed with water before incubating at 60°C for 20 minutes in methanol. After stepwise re-hydration roots were incubated for 25 minutes in 1% periodic acid solution. After washing with water seedlings were incubated in 50 mM Na_2_SO_3_/150 mM HCl solution containing 20 mg/l propidium iodide for 1 hour. After washing with water, roots were further incubated in chloral-hydrate solution (2g chloral-hydrate and 0.5g Glycerol in 1 ml of water) for 30 minutes and mounted on microscopic slide with a 300 μm spacer to avoid root damage. Stacked images from stained roots were recorded using a confocal laser scanning microscope (ZEISS LSM 510 DUO) with a LD LCI-Plan-Apochromat 25×/0.8 DIC Imm Korr objective with glycerol as immersion medium. Excitation wavelength was 561 nm and emission was detected with a long pass filter (LP585). Serial optical sections were reconstituted into 3D image stacks to a depth of 290 μm with in-plane voxel extents of 0.15 μm and a section spacing of 0.9 μm in the z-direction.

### Image processing and analysis

Images were converted to hdf5 format and then stitched to a length of 900–1200 μm from the center of the distal part of the quiescent center (dQC) using xuvTools (Emmenlauer et al., 2009). Thereafter roots were segmented, layers were manually corrected, iRoCs were attached, masks were converted to markers and cell features were extracted to csv/xls files.

## Results

Compared to C3 plants, the increased photosynthetic capacity of C4 plants leads to a more densely branched root system (Wade et al., 2020). As C4 photosynthesis has evolved independently multiple times, it is possible that this increased branching density has a range of molecular foundations. For example, unlike many C3 plants, lateral root growth in *Setaria viridis* does not require a functional AUX1 auxin influx carrier (Huang et al., 2017a; Zhao et al., 2015; Bennett et al, 1996). However, other than this example, little is known about the role of AUX1 during lateral root formation in C4 plants. To test whether the loss of a role for auxin influx has driven increased lateral root density in C4 plants, we generated *aux1* loss-of-function plants in the closely-related domestic plant *Setaria italica* (*siaux1*).

### Setaria aux1 mutants show altered panicle and root development

A *Setaria italica aux1* mutant (*siaux1-1*) was isolated from an ethyl methane sulfonate (EMS)-generated mutant library constructed from wild type *Setaria italica* (Yugu1) parent plants: a drought-resistant and fully sequenced modern cultivar (Bennetzen et al, 2012). Wild-type and *siaux1-1* plants were similar in stature (Fig.1A), but with clear differences in inflorescence development, including fewer primary branches (Fig.1B and Fig.1C), fewer secondary branches and a sparsely branched panicle (Fig.1D-1I). Visualization of panicle structure using scanning electron microscopy indicated that fewer primordia initiated in *siaux1-1* when compared to wild type; those primary branch primordia that did develop were unequally sized at the young panicle stage (Fig.1J-1M). Fewer lateral root primordia were initiated in *siaux1-1* (Fig. 1N and 1O), implying that panicle development and root development are both regulated by *SiAUX1*.

**Figure 1.**
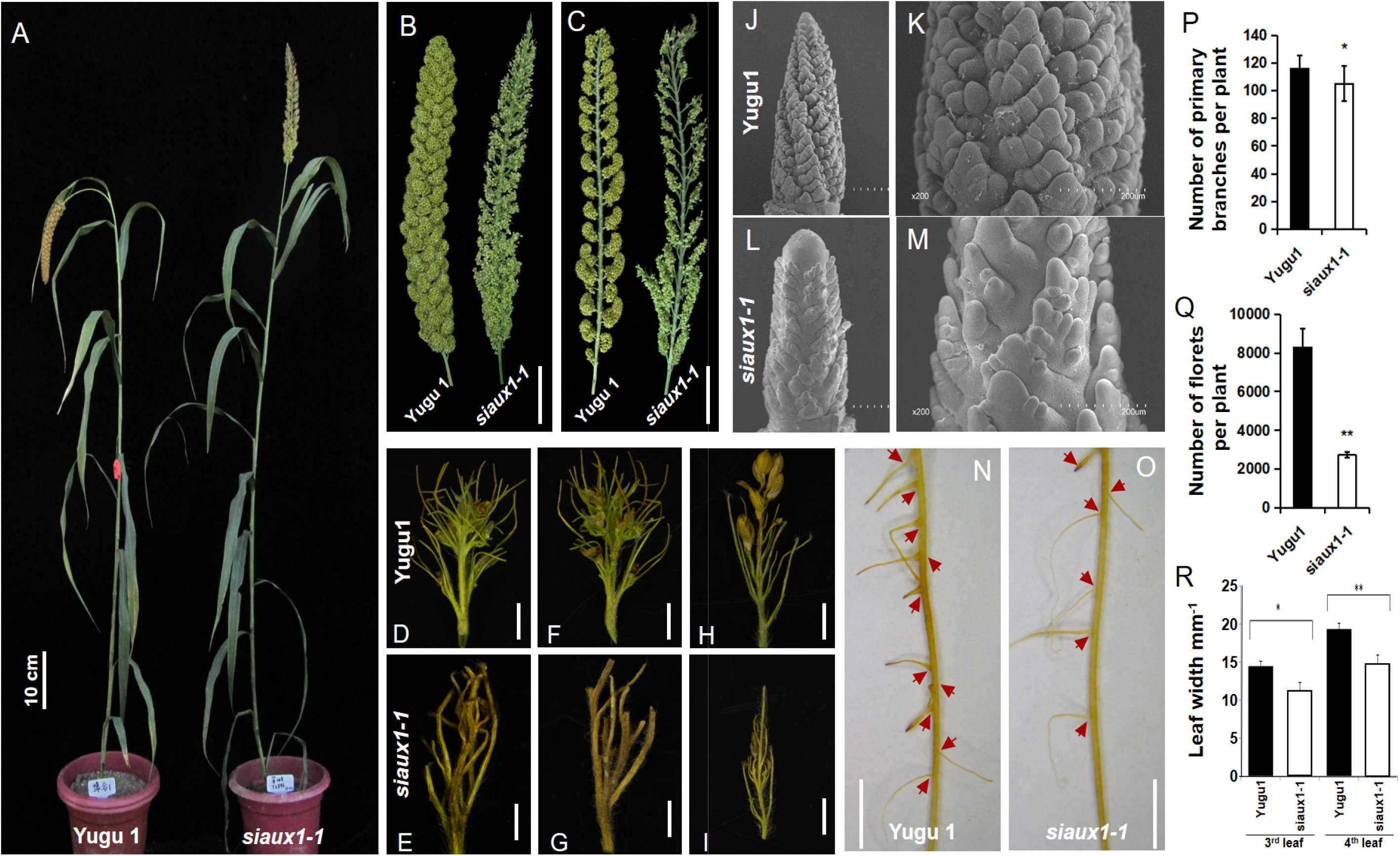
The *Setaria aux1* mutant shows abnormal panicle and root development phenotypes. (A) Whole-plant statue of Yugu1 (wild-type) and *siaux1-1* (mutant). Scale bar = 10 cm. (B) Comparison of panicles of Yugu1 and *siaux1-1* suggests that mutant had sparse panicle. Scale bar = 5 cm. (C) Longitudinal section of Yugu1 and *siaux1-1* panicles clearly demonstrates that mutant had less and abnormal primary branches. Scale bar = 5 cm. (D-I) Florets branches at base (D and E), middle (F and G) and top (H and I) of a panicle. Number of florets branches is significantly reduced in siaux1-1. Scale bar = 2 mm. (J-M) Scanning electron microscopy analysis of Yugu1 (J and K), and siaux1-1 (L and M) young panicles. Less and different primary branch primordia can be observed in mutant. The magnification of the young panicle is 65X with a 250μm scale bar (J and L), and 200X with a 200μm scale bar (K and M). (N-O) Comparison of root phenotype between Yugu1 and *siaux1-1*. Scale bar = 2 cm. (P-R) Quantification of panicle and root phenotypes. Five seedlings were assessed for number of primary branches (P), number of florets per plant (Q), and number of lateral roots per centimeter (R). Asterisks indicate statistically significant differences compared with Yugu1 (*P < 0.05, **P < 0.01; Student’s t test).

In Arabidopsis, LAX2 (Like AUX1) contributes to vascular cell development and vein patterning (Ugartechea-Chirino et al., 2010). *Lax2* (and to some extent also *lax3*) leaves have significantly different vascular structures when compared to wild type, both in their branching pattern and overall symmetry. Furthermore, lateral cross sections in Arabidopsis revealed that stems of *lax2* plants contained a proportionately higher area of vasculature than wild-type plants (Moreno-Piovano et al, 2017), a feature which also drove increased seed yield (Cabello & Chan, 2019). As auxin is generally necessary for the high vein densities that support photosynthesis in leaves of C4 plants (Huang et al, 2017b), we next asked whether *SiAUX1-1* was necessary for plant stature in *Setaria*. In this respect, significant changes in primary branch number, floret number and leaf width were induced by the *siaux1-1* mutation when compared to wild type plants (Fig. 1P-1R).

### SiAUX1 is responsible for a general aux1 phenotype

To identify the responsible *siaux1-1* allele, we constructed three mapping populations including a ‘*siaux1-1*’ × ‘SSR41’ F_2_ population for position cloning and ‘*siaux1-1*’ x ‘Yugu1’ and ‘siaux1-2’ x ‘Yugu1’ BC_1_F_2_ population for Mutmap sequencing. Genetic analyses showed that the *Setaria aux1* phenotype was controlled by a single recessive nuclear gene (Supplementary Table S1). Map-based cloning delimited the *SiAUX1* locus to within a 602-kb region between the molecular markers In5-4153 and In5-4214 on chromosome 5 (Fig. 2A). Mutmap sequencing revealed the presence of only two homozygous SNPs located in the candidate region. One resulted in a premature stop codon within *Seita.5G387200*, which encodes the AUX1 auxin transporter protein (Fig. 2B). The other was a guanine to adenine substitution in an intron of *Seita.5G384900*, with no predicted effects on gene structure. This result indicated that *Seita.5G387200* was the casual gene.

**Figure 2.**
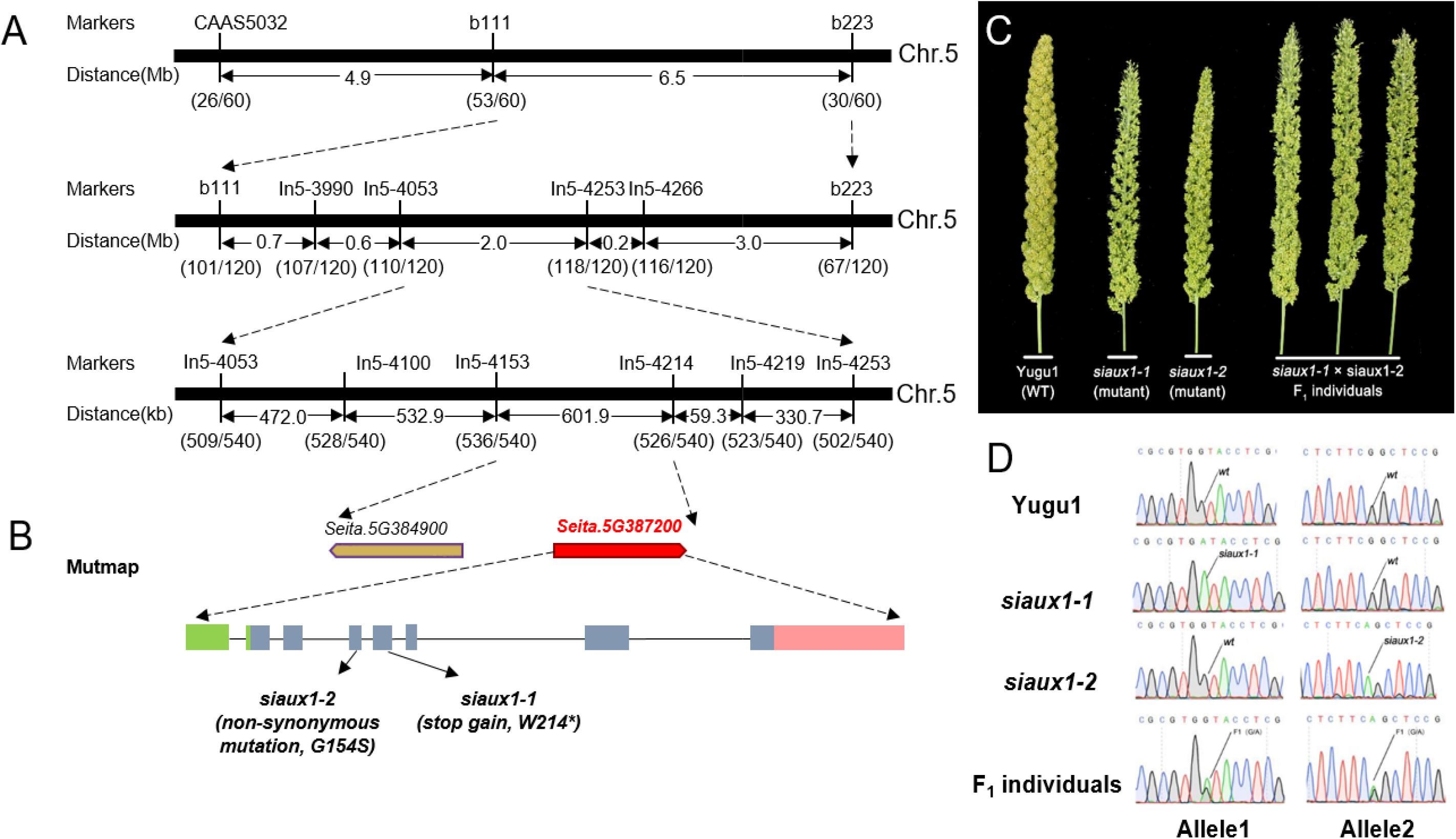
Positional cloning and complementation of *siaux1-1*. (A and B) Mapping the *SiAUX1* locus (Seita.5G387200) using map-based cloning (A) and Mutmap sequencing (B). (C and D) Complementation testing of *Setaria aux1* mutants. Phenotype of panicles of Yugu1, *siaux1-1*, *siaux1-2*, and F_1_ individuals which were generated from *siaux1-1*×*siaux1-2* (C). Sanger sequencing validated SNP variations in each plant.

To further validate whether *Seita.5G387200* was responsible for the phenotype, we re-screened the EMS-treated Yugu1 mutant library, and identified another sparse panicle mutant ‘*siaux1-2’* which had a similar phenotype to *siaux1-1*. We also employed the mutmap strategy to sequence the *siaux1-2* BC_1_F_2_ DNA pool. High-throughput sequencing showed that, genome-wide, *Seita.5G387200*, was the only gene with homozygous non-synonymous mutations in both *siaux1-1* and *siaux1-2*. We therefore conclude that *Seita.5G387200* (hereafter named *SiAUX1*) is responsible for the *Setaria* mutants’ phenotype (Fig. 2B, Supplementary Table S2). To confirm *siaux1-1* and *siaux1-2* were two alleles of *SiAUX1*, we crossed ‘*siaux1-1*‘ and ‘*siaux1-2*‘ plants, and performed complementation testing. The F_1_ hybrid between ‘*siaux1-1*‘ and ‘*siaux1-2*‘ showed an *aux1*-like phenotype, demonstrating both mutations occur in the same gene, *SiAUX1* (Fig. 2C-D). These results confirmed that *SiAUX1* is the causal gene for the *Setaria aux1* phenotype.

### SiAUX1 functions during lateral root emerge and response to gravitropism

In Arabidopsis and rice, AUX1 promotes lateral root primordium development by facilitating the unloading of the most abundant natural auxin, indole-3-acetic acid (IAA), from the vascular transport system at the site of primordium synthesis, causing fewer lateral root primordia to be present in knock-out genotypes (Marchant et al, 2002; Zhao et al, 2015). We therefore generated two independent *SiAUX1*-knock-out lines in foxtail millet using the CRISPR/Cas9 approach (Fig. 3A-D). These knock-out lines showed similar panicle development defects to *siaux1-1* EMS mutants (Fig. 3A-C). In addition, plants overexpressing *SiAUX1* showed more lateral roots when compared with wild-type plants (Fig. 3E). We next scored lateral root density in developing *Setaria* seedlings, comparing wild-type and *siaux1-1* loss-of-function genotypes. The rate of main root growth was not significantly different between wild-type and *siaux1-1* plants (Fig.4A). In wild-type Yugu1 plants, lateral roots began to emerge from the main root axis at five days after germination (DAG). However, *siaux1-1* plants displayed significantly fewer roots than wild type and with an increased incident angle when compared to wild type roots (Fig. 4B-C). In order to ascertain whether SiAUX1 acts during lateral root development in a manner similar to AtAUX1 (initiation) or AtLAX3 (outgrowth) in primordium development, we stained *Setaria* roots to see whether *Siaux1-1* formed the same number of unemerged lateral root primordia as wild-type. In this case, lateral root primordia were seen at three DAG, but no significant difference in their frequency was observed between *Siaux1-1* and wild-type plants (Figure 4D). Our data therefore support the conclusion that SiAUX1 is necessary for lateral root emergence rather than lateral root primordium formation, suggesting a role for SiAUX1 more similar to AtLAX3 than to AtAUX1.

**Figure 3.**
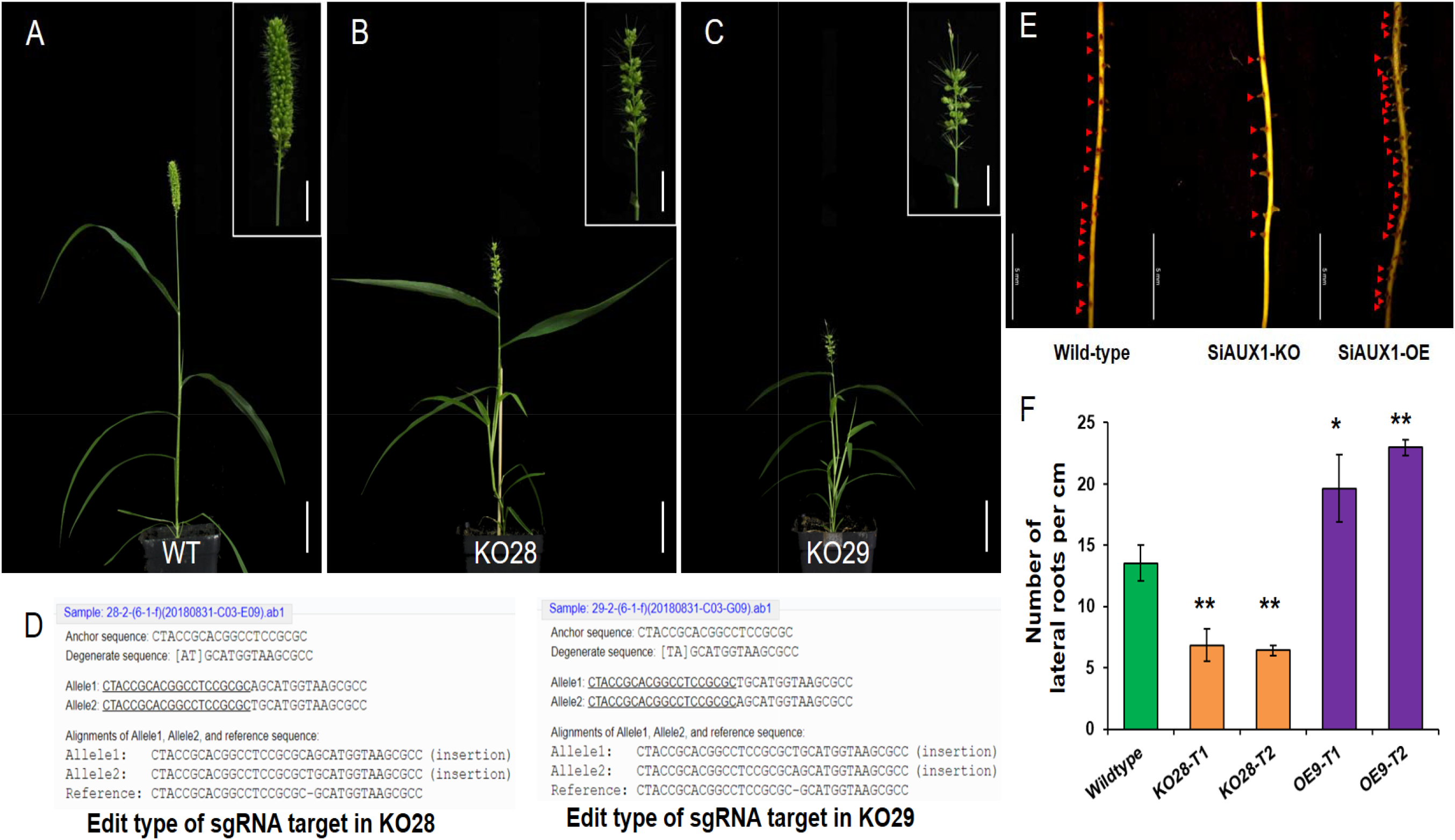
Verification of SiAU.X1 function by CRISPR/Cas9 system in *Setaria*. (A-C) Compared with wild-type plant (A), SiAUX1 knock-out (KO) lines (KO 28 and KO 29) showed sparse panicle phenotype (B and C). Scale bar = 5 cm for whole plant. Scale bar = 2 cm for panicle enlarged. (D) Analysis of CRISPR/Cas9 edit type for positive transgenic lines. (E) Comparison of root phenotype in wild-type, SiAUX1 knock-out lines, and SiAUX1 over-expressed lines. Scale bar = 5 mm. (F) Quantification of root density in independent T_1_ and T_2_ SiAUX1 transgenic lines. Asterisks indicate statistically significant differences compared with wild-type plants (*P < 0.01, **P < 0.001; Student’s t test).

**Figure 4.**
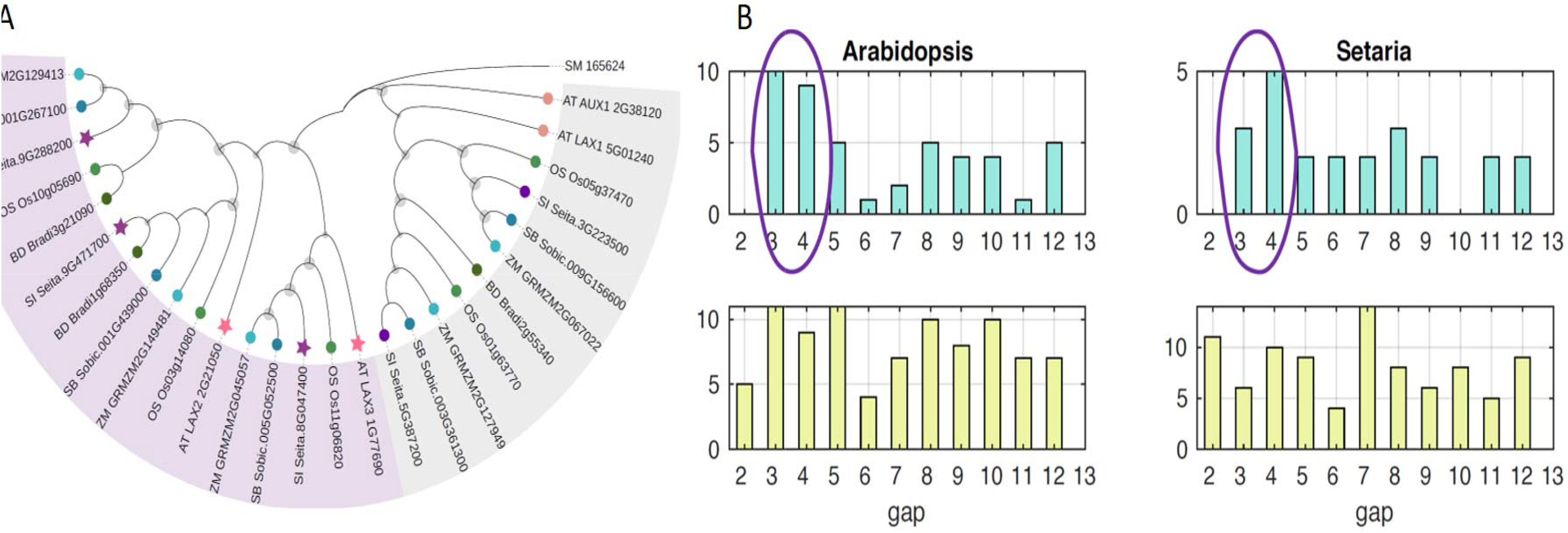
Phylogenetic relationships among *AUX/LAX* family genes (A) Phylogenetic tree for homologs of the *AUX/LAX* family. Pink color highlights clades containing primary auxin responsive genes in Arabidopsis. (B) Grey color marks the remaining clades. Stars marks genes which were taken for motif analysis. Distribution of distances between AuxRE motifs. X-axis – the distance between motifs, Y-axis – the number of motifs found.

The main root of Arabidopsis *aux1* plants display agravitropic growth (Bennett et al, 1996; Marchant et al, 1999), and this phenotype is also seen in *S. italica*. Although the growth rate of *siaux1* plants was indistinguishable from wild-type, growth was plagiotropic, and unresponsive to changes in the gravity vector. Gravitropism differences could not be attributed to differences in the structure of columella cells (the site of gravity perception in the roots), which accumulated starch-containing amyloplasts in the same way as wild-type plants (Fig. 4E).

In Arabidopsis as well as grasses, the expression of AUX1 in non-root hair epidermal cells is required for maintaining rates of root hair outgrowth (Jones et al, 2009). Auxin influx, and particularly its interaction with ethylene-dependent signalling cascades, is also necessary for environmentally responsive root hair growth; for example with respect to low phosphate concentrations (Bhosale et al, 2018; Giri et al, 2018; Rahman et al, 2002). As in Arabidopsis, root hair growth in Yugu1 was stimulated when plants were grown on a reduced-phosphate medium. This response was severely attenuated in *siaux1-1* (Fig. 4F).

### Phylogeny of AUX1 in Setaria italica

The Poaceae (true grasses) are divided into two clades: the PACMAD clade which contains the Paniceae (of which *Setaria* is a member) and the Andropogoneae (containing *Zea*, *Sorghum* and *Miscanthus*), and the BOP clade which contains the Oryzoideaea (Oryza) and the Pooideae (cereals and many pasture grasses).

In Arabidopsis, all four genes of the *AUX1* family are likely to encode functional auxin influx transporters (Peret et al, 2012). There are five homologs of *AUX1\LAX* genes in the genome of *Setaria italica* (*SiAUX1*) and they encode proteins of between 482 and 541 amino acids in length: Seita.5G387200 itself, the homolog of *AtAUX1*, as well as Seita.3G223500, Seita.9G288200 and Seita.9G471700 and Seita.8G047400. Similar domain structures are predicted for all protein products, with most of the differences among sequences being accounted for by highly dissimilar N- and C-termini. Differing codon usage bias between the dicotyledons and grasses hides the evolutionary relationships between homologs at the nucleotide level; therefore, phylogenetic trees were constructed on amino acid sequences.

In order to investigate the phylogenetic structure of the *AUX1\LAX* gene family in the Poaceae, we analysed homologous sequences of five representative species: *Setaria italica*, *Zea mays*, *Sorghum bicolor* from the PACMAD clade, and *Brachypodium distachyon*, and *Oryza sativa* from the BOP clade (Fig. 5A). Our analysis also included the sequences of four Arabidopsis homologs and an outgroup sequence from *Selaginella Moellendorffii*. The outgroup sequence divides all homologs into two clusters: one for *AUX1/LAX1* genes and one for *LAX2/3* genes (Parry et al, 2001). Poaceae homologs formed five distinct clades, each resembling known phylogeny of grasses. The *AUX1/LAX1* cluster comprises two clades of grass homologs. One of the clade corresponds to the *AtAUX1* gene, while the nature of the second clade based on the phylogenetic tree is not clear (Fig. 5A). In the LAX2/3 cluster, two homologs of an At*LAX2* gene were identified in Poaceae genomes, with both PACMAD and BOP clades represented in each. *LAX3* encoded a single homologous Poaceae gene; and in the most current versions of the *Brachypodium distachyon* genome, *LAX3* is not represented.

**Figure 5.**
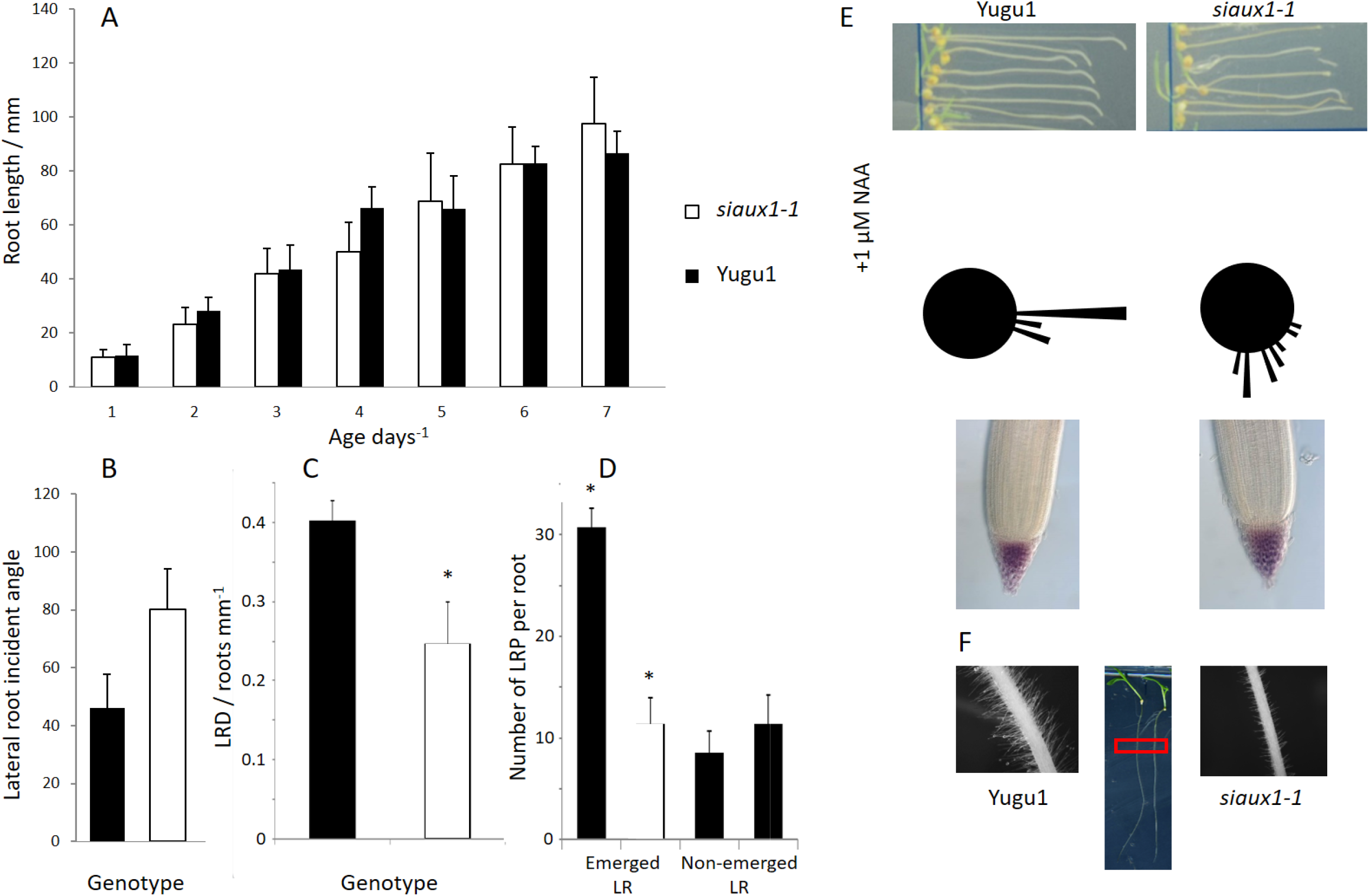
Characterization of the *siaux1-1* root phenotype (A) Growth rate of wild type Yugu1 and *siaux1-1* roots (B) Incidental angle of lateral root growth at 5 days after germination (C) Density of emerged and unemerged lateral root primordia at 5 days after germination (D) Gravitropic response of Yugu1 and *siaux1-1* roots after 90 degree rotation for 23 hours (upper picture) and in the presence of 100 nM NAA (lower picture). Histograms of the gravity response and lugol staining of columella cells. (E) Root hair growth of Yugu1 and *siaux1-1* in phosphate-depleted growth medium

### Promoter structure among auxin-responsive genes in Setaria

In order to examine the basis for auxin-relevant transcriptional responses in Arabidopsis and *Setaria*, we next took primary auxin response genes in Arabidopsis, and asked whether the same one-to-one match that was seen for the AUX/LAX family was present when they were used to search the *Setaria italica* genome. In total, 44 genes whose transcription is rapidly up-regulated by auxin were taken from seven families: *AUX/LAX*, *Aux/IAA*, *GH3*, *PIN*, *ARF*, *LBD* and *SAUR* (Paponov et al, 2008). Unlike for *AUX/LAX* genes, it was not possible to identify direct homologues in the *Setaria* and Arabidopsis genomes for the majority of primary auxin response genes (Figs S1-3). In each gene family, however, we found that auxin-regulated genes in Arabidopsis tended to cluster into clades, helping us to identify the most probable homologs of these genes in *Setaria* (Fig. 5, Figs S1-3).

In order to test whether the *S. italica* genes which were most similar to primary auxin response genes in Arabidopsis could be regulated in a similar fashion, we analysed the frequency of auxin response elements (AuxRE) (TGTCNN) within a region 1kb upstream of their start codon for all *S. italica* and Arabidopsis homologs (Mironova et al, 2014). Auxin response elements bind ARF transcription factors and are a feature of the rapid canonical auxin response (Ballas et al, 1993; Ulmasov et al, 1997). We analyzed the distribution of distances between two ARF-binding motifs spaced no more than 10bp apart in all Arabidopsis and *S. italica* homologs of seven families (*AUX*/*LAX*, *Aux/IAA*, *GH3*, *PIN*, *ARF*, *LBD* and *SAUR)*. Although the overall number of auxin responsive elements did not vary significantly between Arabidopsis auxin responsive and non-responsive genes within the families tested, the distances at which they were spaced were significantly different. AuxREs in the primary auxin responsive genes of Arabidopsis were most frequently separated by 3-4 nucleotides, whereas the distribution for non-regulated genes is almost uniform. When the same families of genes were analyses in *S. italica*, we observed the same partition of *S. italica* homologs, with a separation of 3-4 nucleotides occurring in promoters of genes that were most similar to the primary auxin response genes of Arabidopsis (Fig. 5B). These data are consistent with the hypothesis that clades of auxin responsive genes are highly conserved between dicot plants and grasses.

### Expression profiles of Setaria genes during lateral root emergence

*Siaux1* loss-of-function plants displayed distinctive phenotypic traits both above and below the ground. Leaves were significantly narrower, and lateral root meristems were significantly sparser in knockout plants when compared to Yugu1 (Fig. 1N,O and R). In order to garner an overall view of the molecular mechanisms which could be supporting SiAUX1-dependent growth in the developing seedling, transcription was compared by sequencing mRNA from *siaux1* and Yugu1 plants both before (at the 1-leaf stage) and during (at the 3-leaf stage) lateral root emergence.

Principal component analysis (PCA) of RNA-seq data revealed transcriptional profiles of technical replicates clustered together (Fig. S4A). To compare SiAUX1-dependent transcription profiles we selected transcripts which i) underwent more than 2-fold regulation in either genotype, ii) showed an adjusted *p*-value of less than 0.05, and showed an absolute z-score higher than 2.0 and considered them to be differentially regulated between Yugu1 and *siaux1* seedlings. In total, 33 transcripts fulfilled these criteria, the transcription of 30 of these was up-regulated in *siaux1*, and the transcription of 3 was down-regulated in *siaux1* (Table 1a). Genes predicted to encode proteins involved in nuclear gene regulation were over-represented among transcripts which were up-regulated in *siaux1*. Notably, also up-regulated in *siaux1* was mRNA encoding an aquaporin homologous to the *PIP2* family of aquaporin genes, proteins involved in lignin modification during root development, an enzyme involved in folate metabolism. Most of the 33 transcripts which were significantly regulated between Yugu1 and *Siaux1* showed a highly dynamic expression profile in roots and shoots during the first days after germination, according to publicly available *S. italica* expression data (Fig. S4B).

**Table 1.**
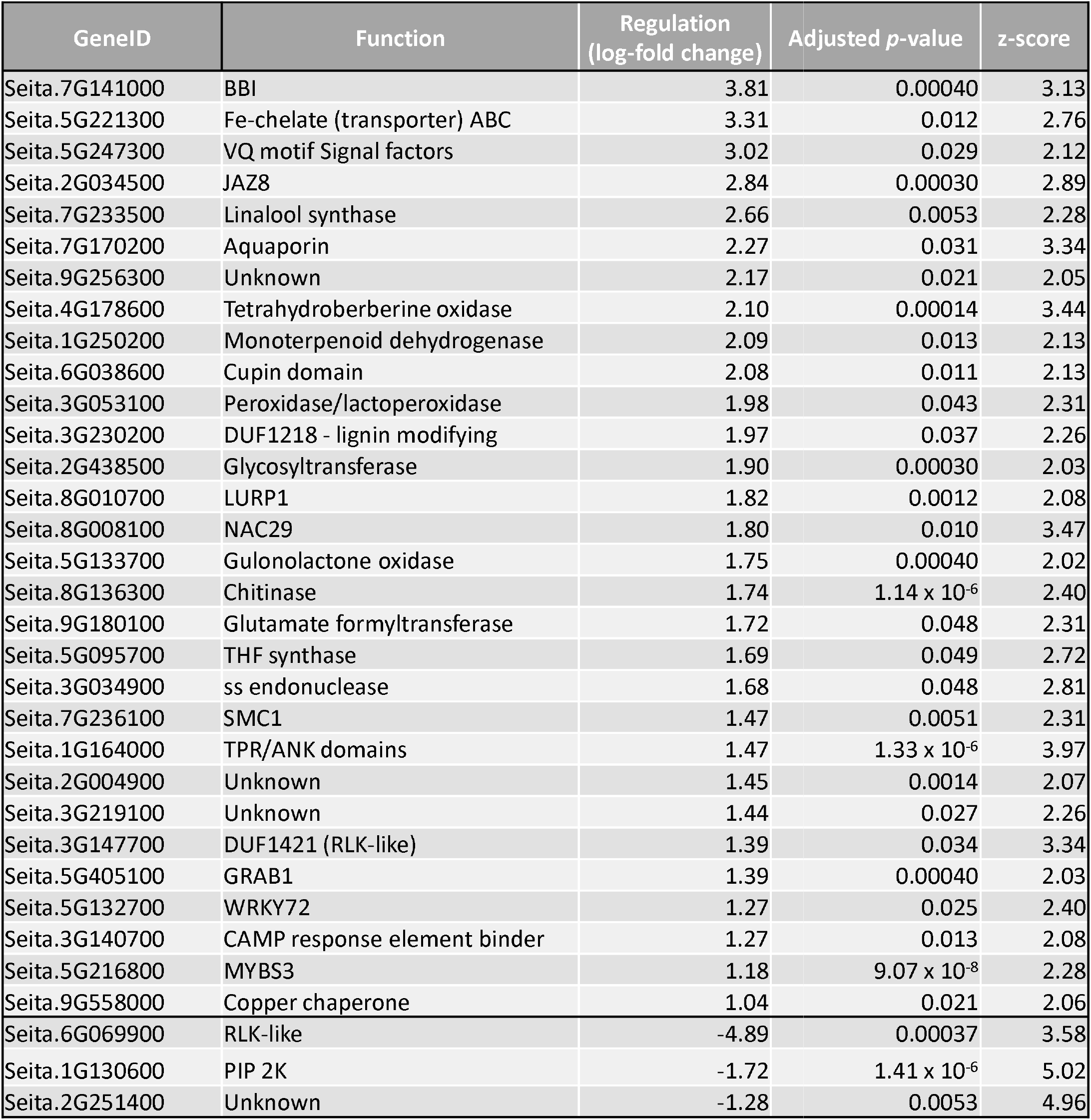
A summary of genes which are differentially regulated in *siaux1-1* during seedling growth in *Setaria italica*. RNAseq was carried out on tissue samples taken from roots and leaves during both one-leaf and three-leaf developmental stages.

| GeneID | Function | Regulation<br>(log-fold change) | Adjusted *p*-value | z-score |
| --- | --- | --- | --- | --- |
| Seita.7G141000 | BBI | 3.81 | 0.00040 | 3.13 |
| Seita.5G221300 | Fe-chelate (transporter) ABC | 3.31 | 0.012 | 2.76 |
| Seita.5G247300 | VQ motif Signal factors | 3.02 | 0.029 | 2.12 |
| Seita.2G034500 | JAZ8 | 2.84 | 0.00030 | 2.89 |
| Seita.7G233500 | Linalool synthase | 2.66 | 0.0053 | 2.28 |
| Seita.7G170200 | Aquaporin | 2.27 | 0.031 | 3.34 |
| Seita.9G256300 | Unknown | 2.17 | 0.021 | 2.05 |
| Seita.4G178600 | Tetrahydroberberine oxidase | 2.10 | 0.00014 | 3.44 |
| Seita.1G250200 | Monoterpenoid dehydrogenase | 2.09 | 0.013 | 2.13 |
| Seita.6G038600 | Cupin domain | 2.08 | 0.011 | 2.13 |
| Seita.3G053100 | Peroxidase/lactoperoxidase | 1.98 | 0.043 | 2.31 |
| Seita.3G230200 | DUF1218 - lignin modifying | 1.97 | 0.037 | 2.26 |
| Seita.2G438500 | Glycosyltransferase | 1.90 | 0.00030 | 2.03 |
| Seita.8G010700 | LURP1 | 1.82 | 0.0012 | 2.08 |
| Seita.8G008100 | NAC29 | 1.80 | 0.010 | 3.47 |
| Seita.5G133700 | Gulonolactone oxidase | 1.75 | 0.00040 | 2.02 |
| Seita.8G136300 | Chitinase | 1.74 | 1.14 x 10<sup>-6</sup> | 2.40 |
| Seita.9G180100 | Glutamate formyltransferase | 1.72 | 0.048 | 2.31 |
| Seita.5G095700 | THF synthase | 1.69 | 0.049 | 2.72 |
| Seita.3G034900 | ss endonuclease | 1.68 | 0.048 | 2.81 |
| Seita.7G236100 | SMC1 | 1.47 | 0.0051 | 2.31 |
| Seita.1G164000 | TPR/ANK domains | 1.47 | 1.33 x 10<sup>-6</sup> | 3.97 |
| Seita.2G004900 | Unknown | 1.45 | 0.0014 | 2.07 |
| Seita.3G219100 | Unknown | 1.44 | 0.027 | 2.26 |
| Seita.3G147700 | DUF1421 (RLK-like) | 1.39 | 0.034 | 3.34 |
| Seita.5G405100 | GRAB1 | 1.39 | 0.00040 | 2.03 |
| Seita.5G132700 | WRKY72 | 1.27 | 0.025 | 2.40 |
| Seita.3G140700 | CAMP response element binder | 1.27 | 0.013 | 2.08 |
| Seita.5G216800 | MYBS3 | 1.18 | 9.07 x 10<sup>-8</sup> | 2.28 |
| Seita.9G558000 | Copper chaperone | 1.04 | 0.021 | 2.06 |
| Seita.6G069900 | RLK-like | -4.89 | 0.00037 | 3.58 |
| Seita.1G130600 | PIP 2K | -1.72 | 1.41 x 10<sup>-6</sup> | 5.02 |
| Seita.2G251400 | Unknown | -1.28 | 0.0053 | 4.96 |

### The cellular structure of the siaux1-1 root apical meristem

In Arabidopsis, the distribution of auxin across the root apical meristem (RAM) serves as a positional cue which determines cell identity laterally (Aida et al, 2004), and meristem size longitudinally (Dello Ioio et al, 2008). Auxin also contributes to the specification of specific files of cells as vasculature within the stele (Bishopp et al, 2011a). Models have predicted that both cellular auxin influx and efflux both contribute to the sites of auxin accumulation in the RAM (Band et al, 2014; Moore et al, 2017). Auxin influx carriers are known to contribute to the control of RAM structure as, in Arabidopsis, *aux1* roots have longer epidermal cells (Strader et al, 2010), *lax2* plants display quiescent cell (QC) division (Zhang et al, 2013), and loss of function *aux/lax* genotypes display a generally aberrant root cap and columella organization (Ugartechea-Chirino et al, 2010). Despite these differences, meristem length in *aux1* Arabidopsis plants is indistinguishable from wild type and cell length in the elongation zone remains unaffected (Street et al, 2016). In rice, cell cycle activity in the RAM appears also to be dependent on *OsAUX3*, but in contrast, here loss-of-function mutants displayed a short-root phenotype (Wang et al, 2019).

In order to test whether *SiAUX1* is necessary for the maintenance of meristem structure in *Setaria*, we used iRoCS, a root coordinate system (Schmidt et al, 2014), to label and measure the volumes of each cell of *Siaux1* RAM and compare them to cells from identical positions in wild type roots.

### Root Apical Meristem structure

We first compared the sizes of the division zone (DZ; here defined as the distal region of the root apical meristem which contains relatively small isodiametric cells in its outer layers) in wild type and *aux1* roots by plotting epidermal cell length against longitudinal distance from the quiescent centre (QC). No significant difference was observed in meristem length between *siaux1-1* and wild type roots when either epidermal cell length or absolute deviation in epidermal cell volume was used (Figure S5A-B). Despite there being no difference in DZ length, epidermis cells in the RAM had a significantly larger absolute volume in *siaux1* than wild type roots (Figure S5C). However, consistent with there being no significant difference in DZ length between RAMs of *Siaux1* and wild type plants, the difference in cell volumes between *siaux1* and wild-type plants did not vary with longitudinal distance (Fig S5B-C).

The *Setaria* RAM contains three layers of cortex cells which separate the epidermal and endodermal cell layers (Fig. 6). In order to ascertain whether AUX1 differentially affected cell growth in cell layers of the root apical meristem, we compared relative meristem structures by mapping the frequency distributions of cell volumes between 300μm and 800μm from the QC in the outer five cell layers. As was seen in Arabidopsis, the largest differences were seen when epidermal cell size of the *Siaux1* and wild type RAM was compared. The wild type RAM contained fewer small cells than did the *Siaux1* RAM in a corresponding position. This trend was also observed in other cell layers tested, but was less pronounced in inner cell layers (Fig. 6). We therefore conclude that, as has been described in Arabidopsis (Strader et al, 2010) AUX1 function limits cell size in the DZ. Although this effect is less pronounced in inner cell layers, as has also been observed in Arabidopsis, AUX1 function does not limit RAM size in *Setaria*.

**Figure 6.**
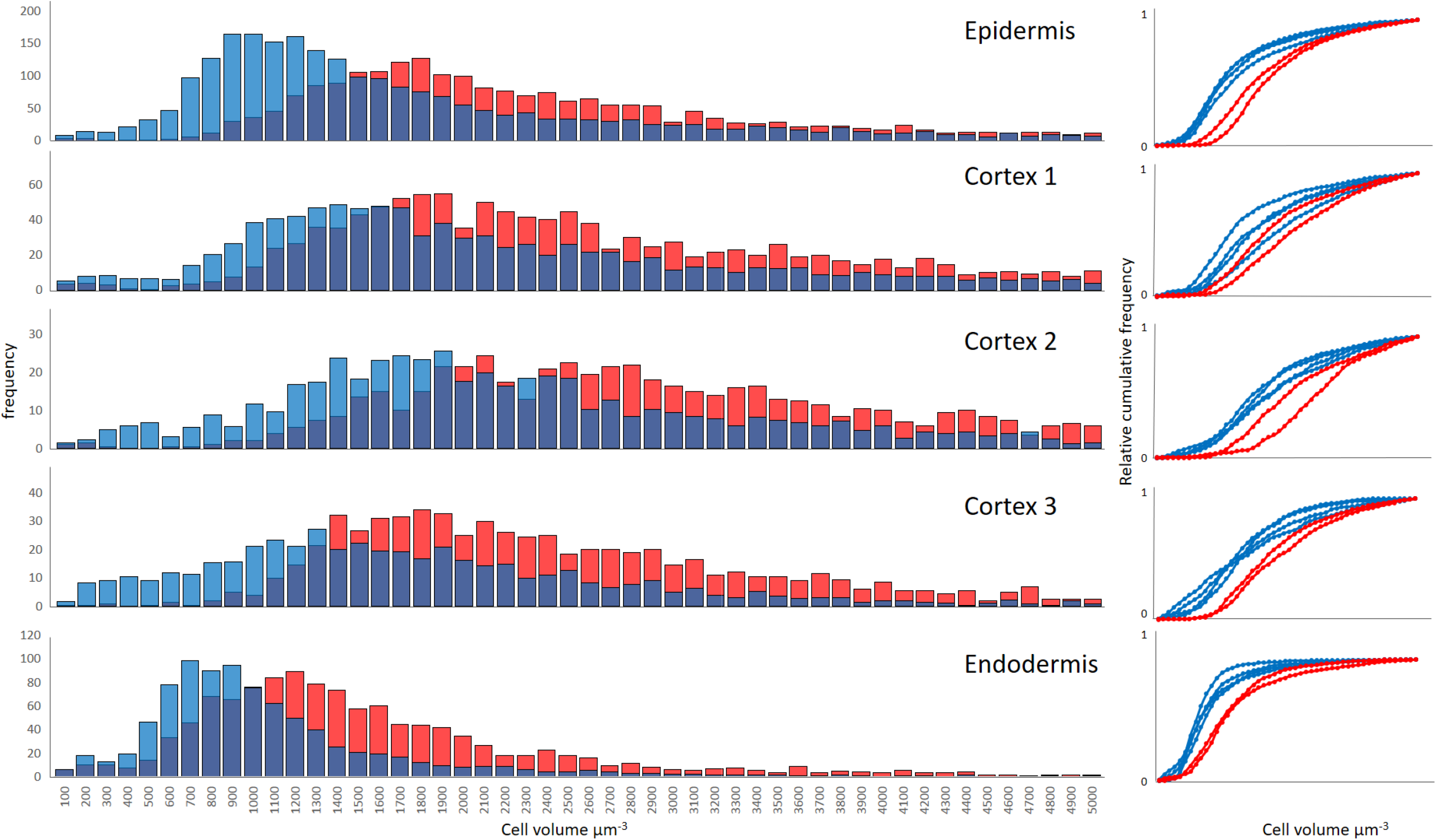
Cell file-specific distribution of cell volumes in Yugu1 and *siaux1-1* root apical metistems Left: cell volumes from the root apical meristem at between 300 and 800 μm from the quiescent center were sorted into bins of 100 μm^3^, and their frequencies plotted. The averages of four Yugu1 roots (blue) and two *siaux1-1* (red) roots are shown Right: cumulative relative frequencies of individual roots

In Arabidopsis roots, AUX1-dependent auxin efflux helps to define the structure of vasculature in developing primary roots by controlling both the pattern of cells within the central stele and the rate of their differentiation (Fabregas et al, 2015). The differences in root architecture between grasses and eudicots are also present at a cellular level within the meristem of the primary root. We were therefore interested in whether AUX1 function was preserved in this different morphological context. In *Setaria* between five and six strands of xylem are present (Sebastian and Dinney, 2017). Our analysis also bore this observation out: of seven segmented wild type roots, six contained six strands of metaxylem at a distance of 250μm proximal to the QC, and one had five. In contrast, all of the four *Siaux1* roots segmented only displayed five metaxylem strands. 3D reconstruction was also able to identify six protophloem strands interspaced between files of metaxylem cells (Fig. 7). We therefore conclude that AUX1-dependent auxin influx is also necessary for vascular patterning in *Setaria*, despite the morphological differences between primary roots of eudicots and panicoid grasses. Vascular structure was reflected in pericycle cell layer: files of larger pericycle cells were found proximal to protophloem strands, and protoxylem cells closer to smaller pericycle cells. The volume of small pericycle cells (proximal to protoxylem cells) was independent of their distance from the QC (up to 800 μm away) and therefore did not elongate in the transition zone (Fig. 7). The spatial relationship between pericycle cell size and vascular structure was preserved in *Siaux1*, which contained only five regions of large pericycle cells in any given radial cross section. The difference in pericycle cell volumes in such a section appeared exaggerated in the *Siaux1* RAM when compared to wild type.

**Figure 7.**
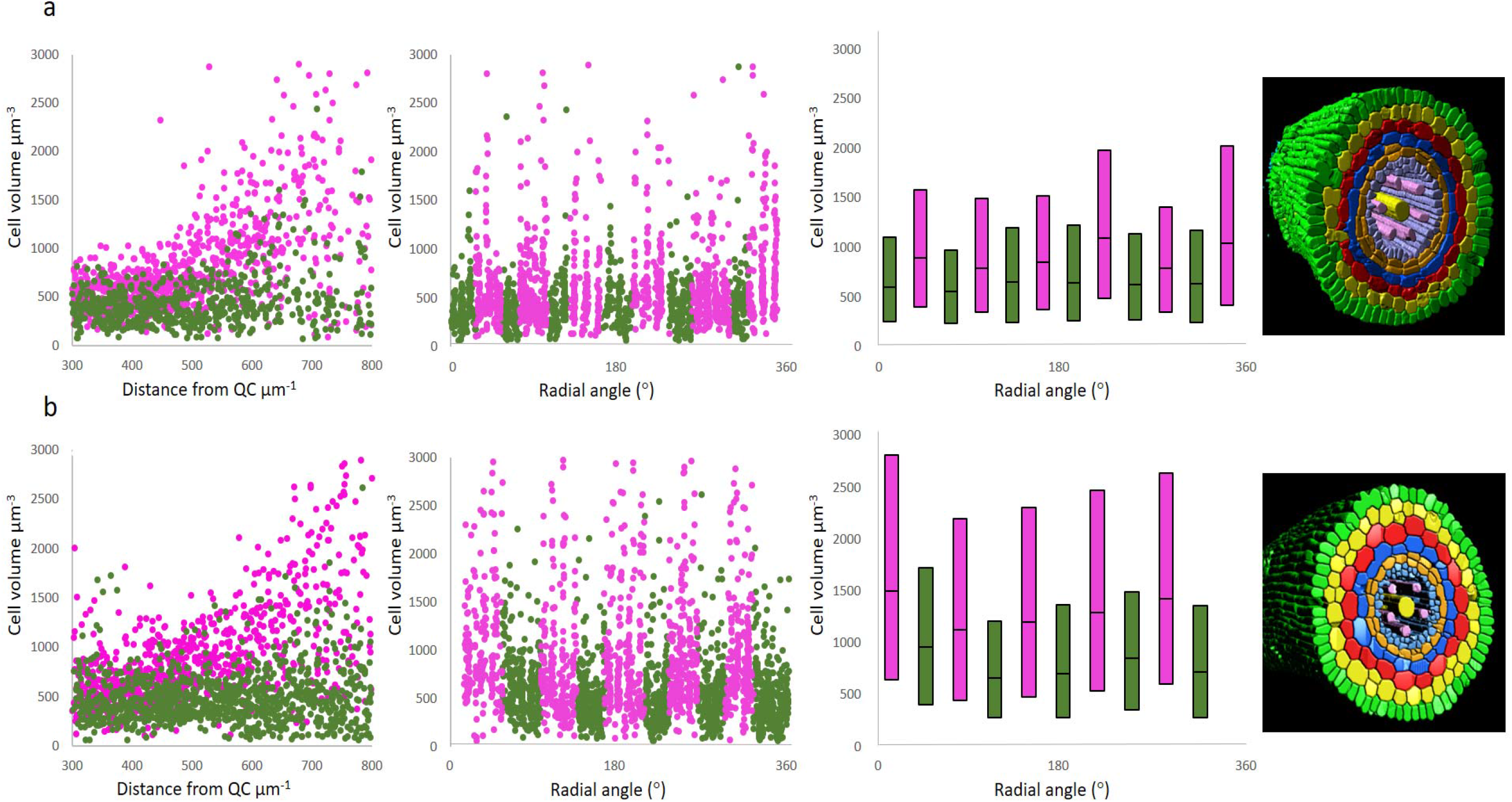
Pericycle cell volume in the *Setaria* RAM. RAM of (a) wild type and (b) *Siaux1-1* loss-of-function mutant plants. Two populations of pericycle cell (pink and dark green) were discernable and reflected directly the subtending vascular structure (right). All cell volumes were pooled between 300 and 800 μm from the QC. Bars indicate lower and upper quartiles and median cell volume values. Radial sections were taken at 250 μm from the QC. Pith cells not shown.

## Discussion

Monocots and dicots plants are estimated to have diverged around 140 million years ago (Chaw et al, 2004). However, our experiments indicate that the function of AUX1 has been largely conserved between classes. There is also convincing evidence that many of these roles are also conserved in maize, another member of the PACMAD clade of panicoid grasses (Huang et al, 2017a).

In Arabidopsis, AUX1- and LAX3-dependent increases in cellular auxin concentration activate developmental programs which lead to lateral root growth. These programs require the auxin-mediated transcription of Lateral Organ Boundaries Domain (LBD)/Asymmetric Leaves2-Like (ASL) proteins (Lee et al, 2015). It is likely that this transcriptional signature in lateral root development is shared in *Setaria* as *LBD16* and *LBD18* genes have homologs in its genome. Furthermore, *Setaria LBD16* and *LBD18* promotor regions (at loci *Seita.1g369600* and *Seita.9g470000* respectively) also contain a spacing pattern of auxin response elements typical of primary auxin response genes (Figure S3). *ARF7* and *ARF19* (not themselves auxin response genes) mediate the AUX1/LAX3-dependent LBD16/18 response, and also have direct homologs in *Setaria* (*Seita.6g212000*, *Seita.4g240600* and *Seita.1g090900*). *LBD29* regulates *LAX3* expression in Arabidopsis and is a primary auxin response gene. However, we do not expect the three genes in *Setaria* which are most similar to *AtLBD29* to also be primary auxin response genes as their promoter regions do not contain a high frequency of auxin response elements (Porco et al, 2016). Our analysis found that the overall number of repeated auxin response elements in a promoter region was a poor predictor of whether the downstream open reading frame is a homolog of a gene that is transcribed upon auxin treatment in Arabidopsis (Paponov et al, 2008). Spacing of auxin response elements is crucial for ARF binding and subsequent gene transcription (Boer et al, 2014); this factor is also likely to be a key determinant of gene activity in *Setaria*. Consistent with this hypothesis is a bias in the spacing of auxin responsive elements in both Arabidopsis and *S. italica* in primary auxin response genes and their homologs (Figure 4b).

The isolation of the *sparse panicle1* (*spp1*) allele in *Setaria viridis*, and its identification as an AUX1 loss-of function mutation has eloquently illustrated the value of using foxtail millet as a model C4 grass (Huang *et al*., 2017a). Unlike in *spp1*, both *Siaux1* and *Ataux1* plants displayed a very similar reduction in lateral root density. In rice, AUX1 contributes to the regulation of lateral root density (Zhao et al, 2015). Together, these observations suggest AUX1 does not function in lateral formation in *S. viridis*. It is now important to test whether this functional specification is characteristic of other C4 species within the Paniceae and Andropogoneae. Richer soils adapted for cultivation during the domestication of *S. italica* may have eased the evolutionary pressure to form a highly branched root system able to support C4 photosynthesis. It is now a high priority to investigate the molecular basis for AUX1-independent lateral root formation in *S. viridis*, and find out whether it is more widespread among C4 species.

In monocots, lateral roots are initiated in phloem pole pericycle and endodermis cells (Bell and McCully, 1970). In monocots plants, as in dicots, auxin plays a central role throughout the early stages of lateral root development, as can be seen in the prominence of auxin signalling mutants with lateral root phenotypes (Yu et al., 2016). In arabidopsis, emerging lateral roots making their way through the cortex and epidermis cell layers with the help of LAX3 and the aquaporin PIP2.1 which serves to weaken the bonds between cell files. Here, the dynamic expression of *PIP2.1* is crucial during lateral root development is necessary for emergence (Péret et al., 2012). In *S. italica*, *Seita.7G170200* is both upregulated in *siaux1* loss-of-function plants and has a highly dynamic expression profile in seedlings (Table 1 and Fig. S4B). Folate biosynthesis interacts with root development to affect the gravitropic response (Nziengui et al., 2018). Tetrahydrofolate synthase (*Seita.5G0995700*) gene expression is also differentially regulated in *siaux1* loss-of-function plants; testing the effect of *para*-amino benzoic acid application on lateral root formation and gravitropism in *S. viridis* could reveal mechanisms for AUX1 functional suppression. A berberine bridge enzyme (BBE)-like enzymes oxidation of aromatic allylic alcohols during lignification in the cell wall (Daniel et al., 2015). Another transcript differentially expressed in *siaux1* loss-of-function seedlings, *Seita.3G230200*, encodes a protein with a characteristic BBE-enzyme domain structure predicted to be involved in lignin modification with a potential role in cell wall dynamics during lateral root emergence. The expression of BBE-enzymes has already been shown to be regulated by an *LBD*-transcription factor complex during root development in Arabidopsis (Xu et al., 2018). Heavy metals such as copper and iron positively influence lateral root growth in a jasmonate-specific manner (Ferrieri et al., 2017). In this context, the products of three *aux1*-regulated genes may be relevant during lateral root formation: *Seita.5G221300* and *Seita.9G558000* in heavy metal ion transport, and their potential interaction with pathways involving the transcriptional regulator *JAZ8* (*Seita.2G034500*).

In order to compare the root phenotype of SiAUX1 and wild type plants, we adapted iRoCS, a root coordinate system which describes the relative position of each cell of the root tip with respect to the central spline of the root. This allows us to remove naturally occurring bends of a few micrometers in the root tip for a direct overlay and comparison of root architecture at a cellular resolution. Our analysis shows that SiAUX1 restricts cell size in the root apical meristem, predominantly in the epidermis (Figure 6). That the epidermis appears to be predominant in the determination of meristem length is surprising, as at least in maize, *AUX1* mRNA expression is confined to endodermis and pericycle cells (Hochholdinger et al, 2000). In Arabidopsis the interaction between the transport of auxin and signalling, in response to both auxin and cytokinin, determines vascular structure. By inhibiting the efficacy of cytokinin or auxin signalling pathways it is possible to alter the relative number of protoxylem cells in the stele (Bishopp et al, 2011a). *Siaux1* were shown, in the current analysis, to sometimes display only five vascular strands instead of the usual six typical of wild-type plants (Figure 7). In Arabidopsis, auxin efflux rather than auxin influx regulates the relative activities of auxin and cytokinin signalling pathways in neighbouring cell files (Bishopp et al, 2011b), whereas auxin efflux and influx routes act in concert to regulate the spacing of lateral roots (Peret et al, 2013). It is therefore possible that the interaction of auxin transport systems is regulated in a different manner in *Setaria*. We suggest here that determining at high genetic and spatial resolution which patterning pathways are functionally equivalent in *Setaria* and Arabidopsis promises a potentially important platform on which our understanding of grass crop species can be built.

## Acknowledgements

The authors would gratefully like to acknowledge the funding of the National Natural Science Foundation of China (31660423, 31871692); the National Key R&D Program of China (nos. 2019YFD1000700 and 2019YFD1000702); the Bundesministerium für Bildung und Forschung (BMBF 031B0556), the Excellence Initiative of the German Federal and State Governments (EXC 294) and German Research Foundation (DFG SFB746, INST 39/839,840,841) and Deutsche Agentur für Luft-und Raumfahrt (DLR 50WB1522). We would also like to acknowledge the expert technical support of Thorsten Falk and Kinan Alzouabi. The authors declare that no competing interests exist.

## Author contributions

ST, planned and performed gene cloning experiments, data analysis, and prepared the manuscript; JX, performed experiments; MS and SA planned and performed experiments, TP made all cell-by-cell root phenotype measurements, AI, GM and MS performed all phylogenetic analyses, FR prepared experimental protocols, HZ, performed field work; YG, performed F_1_ complementation tests; YS and CW prepared transgenic material; GJ and XL helped design experiments; XD, KP and WT directed and guided the program, analyzed data and prepared the manuscript.

## Data availability statement

The data that support the findings of this study are openly available at EMBL-EBI (http://www.ebi.ac.uk/), reference number PRJEB44706.

## Supplementary information

**Supplementary Table S1.** Genetic analysis of *siaux1-1* mapping population.

**Supplementary Table S2.** Mutmap sequencing of *siaux1-1*×Yugu1 and *siaux1-2*×Yugu1 BC_1_F_2_ mapping population.

**Supplementary Table S3.** Primer lists for major experiments

**Supplementary Table S4.** Auxin-related gene homologs in Arabidopsis and *Setaria*

**Supplementary Figure S1** Phylogenetic tree for homologs of auxin signaling genes family. A) *ARF* genes. B) *AUX/IAA* genes. Pink color highlights clades containing primary auxin responsive genes in Arabidopsis. Grey color marks the remaining clades. Stars marks genes which were taken for motif analysis illustrated in figure 4B

**Supplementary Figure S2**

Phylogenetic tree for homologs of auxin responsive genes. A) *PIN* genes. B) *SAUR* genes. Pink color highlights clades containing primary auxin responsive genes in Arabidopsis. Grey color marks the remaining clades. Stars marks genes which were taken for motif analysis illustrated in figure 4B

**Supplementary Figure S3**

Phylogenetic tree for homologs of the *LBD* family. Pink color highlights clades containing primary auxin responsive genes in Arabidopsis. Grey color marks the remaining clades. Stars marks genes which were taken for motif analysis illustrated in figure 4B

**Supplementary Figure S4**

A) Principal component analysis of RNAseq analysis. Samples are labelled according to age 1L (one-leaf stage), 3L (three-leaf stage) and tissue type, root (Rt), Lf (leaf) and sam.pnc (panicle). Tissue was collected from either wild-type (Yu) or *aux1* plants (*aux*). B) Expression analysis of genes differentially regulated between *aux1* and wild-type plants. Data were obtained from the following publically available *Setaria italica* gene expression resource: http://www.setariamodel.cn/#/expression.

**Supplementary Figure S5** Analysis of epidermal cell dimensions in the *siaux1-1* root apical meristem

Epidermal cell volume and length are given as functions as distance from the quiescent center. Average cell lengths and volumes in the epidermis between 100 and 500 μm from the QC. iRoCS is used to superimpose 2 Yugu1 roots (blue) and 2 *siaux1-1* roots (red)

