## Supplemental Figures for "The role of *AUX1* during lateral root development in the domestication of the model C4 grass *Setaria italica*"

### Slide 1
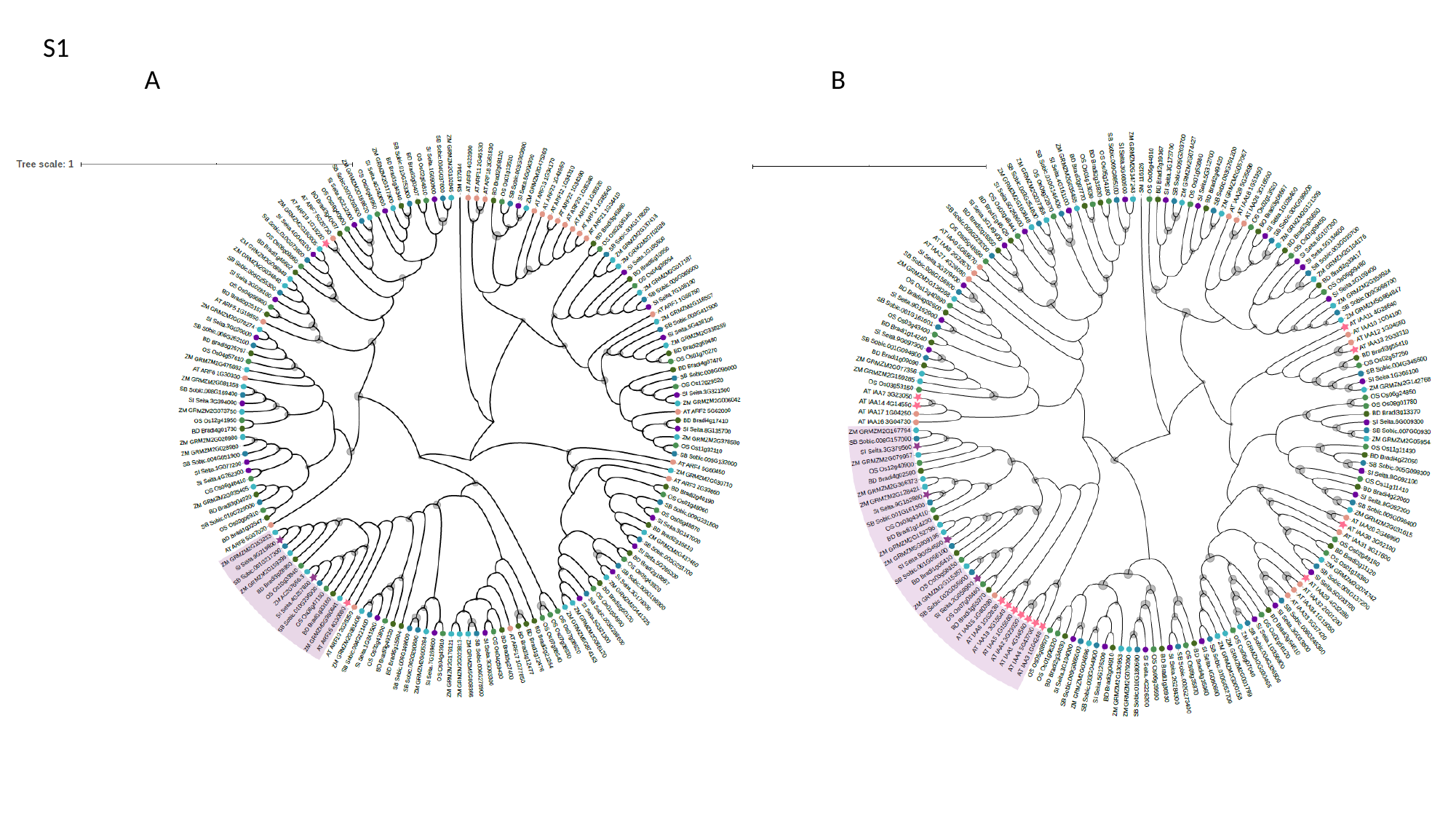

S1
A
B

### Slide 2
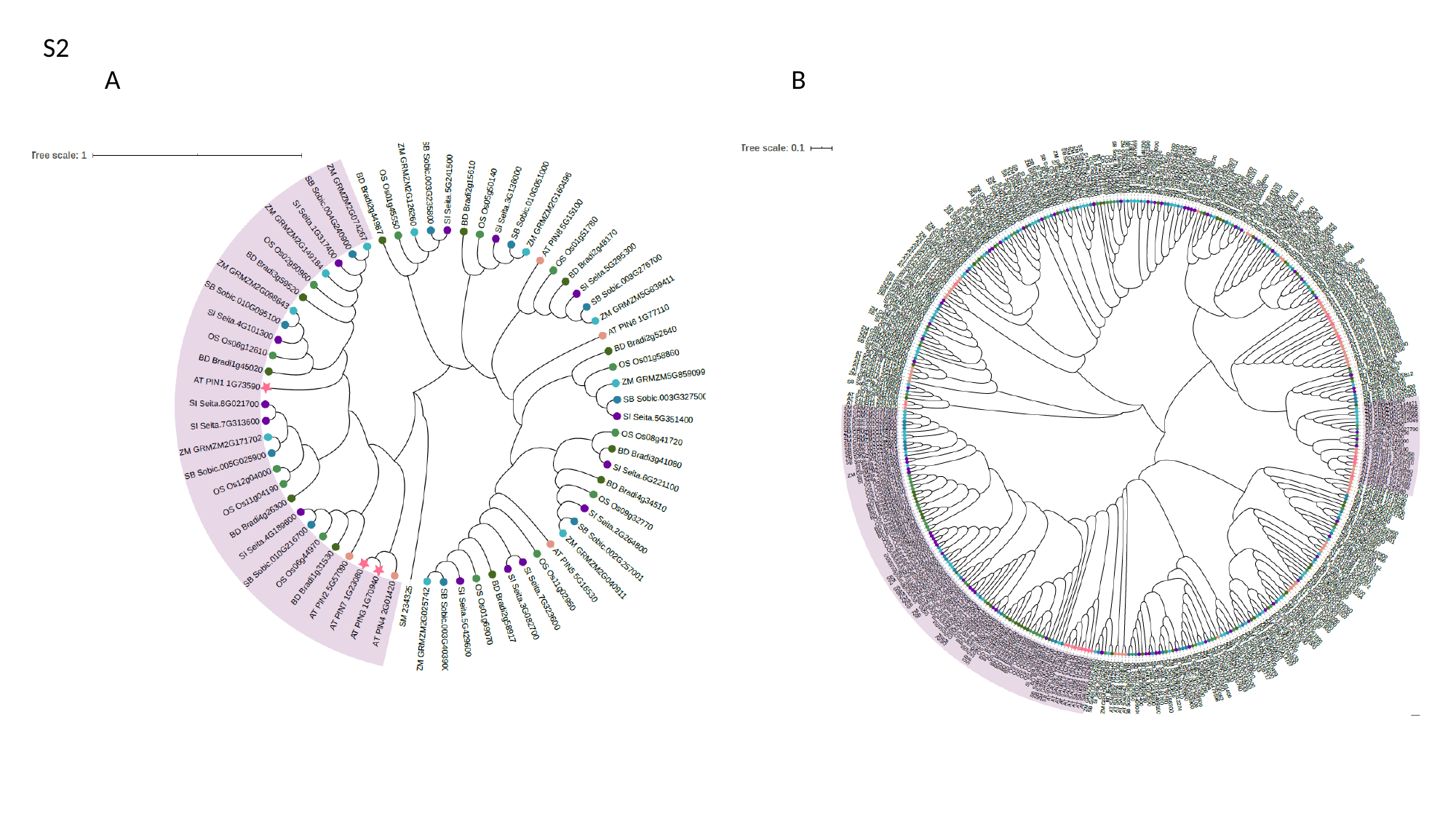

S2
A
B

### Slide 3
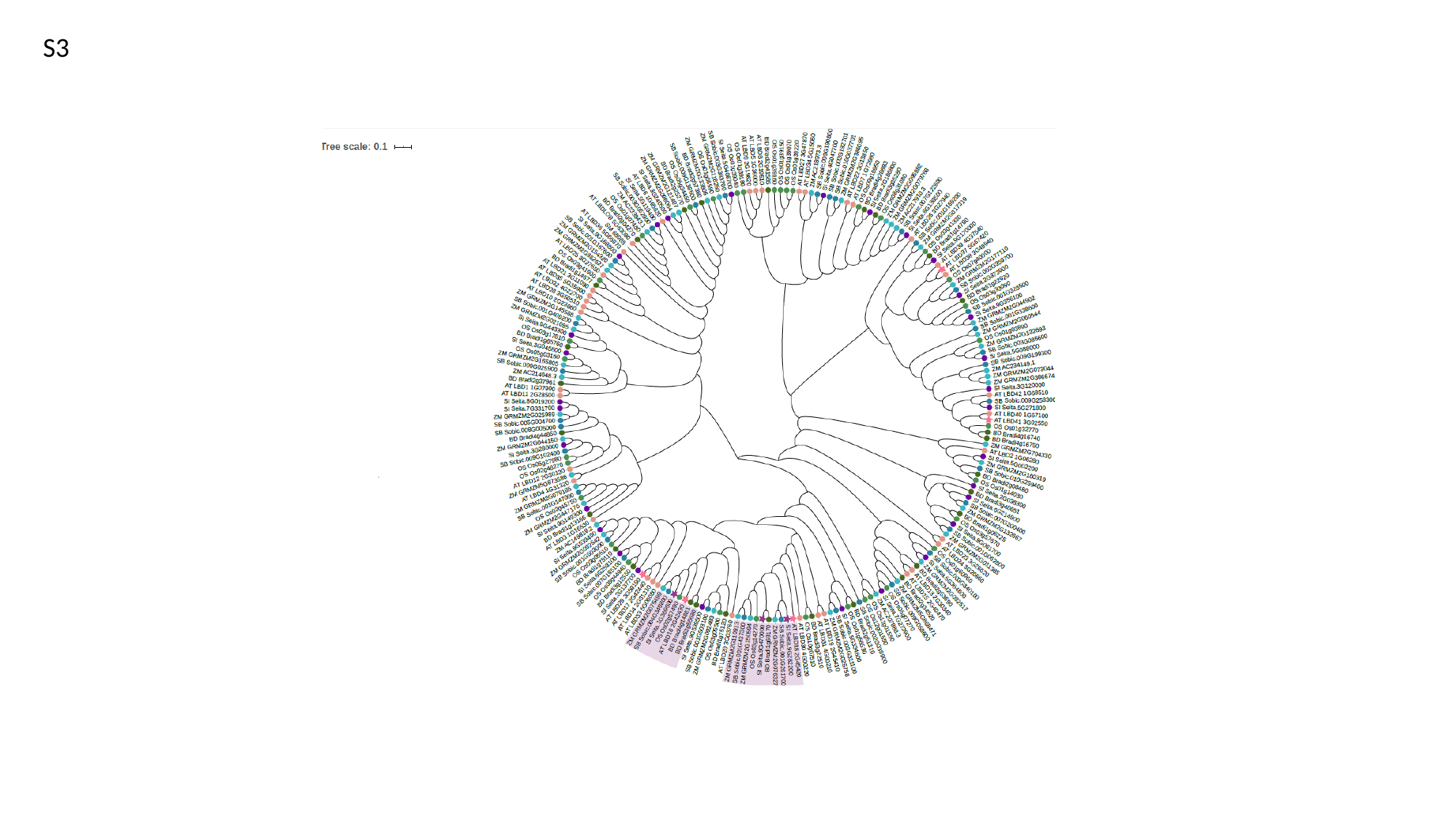

S3

### Slide 4
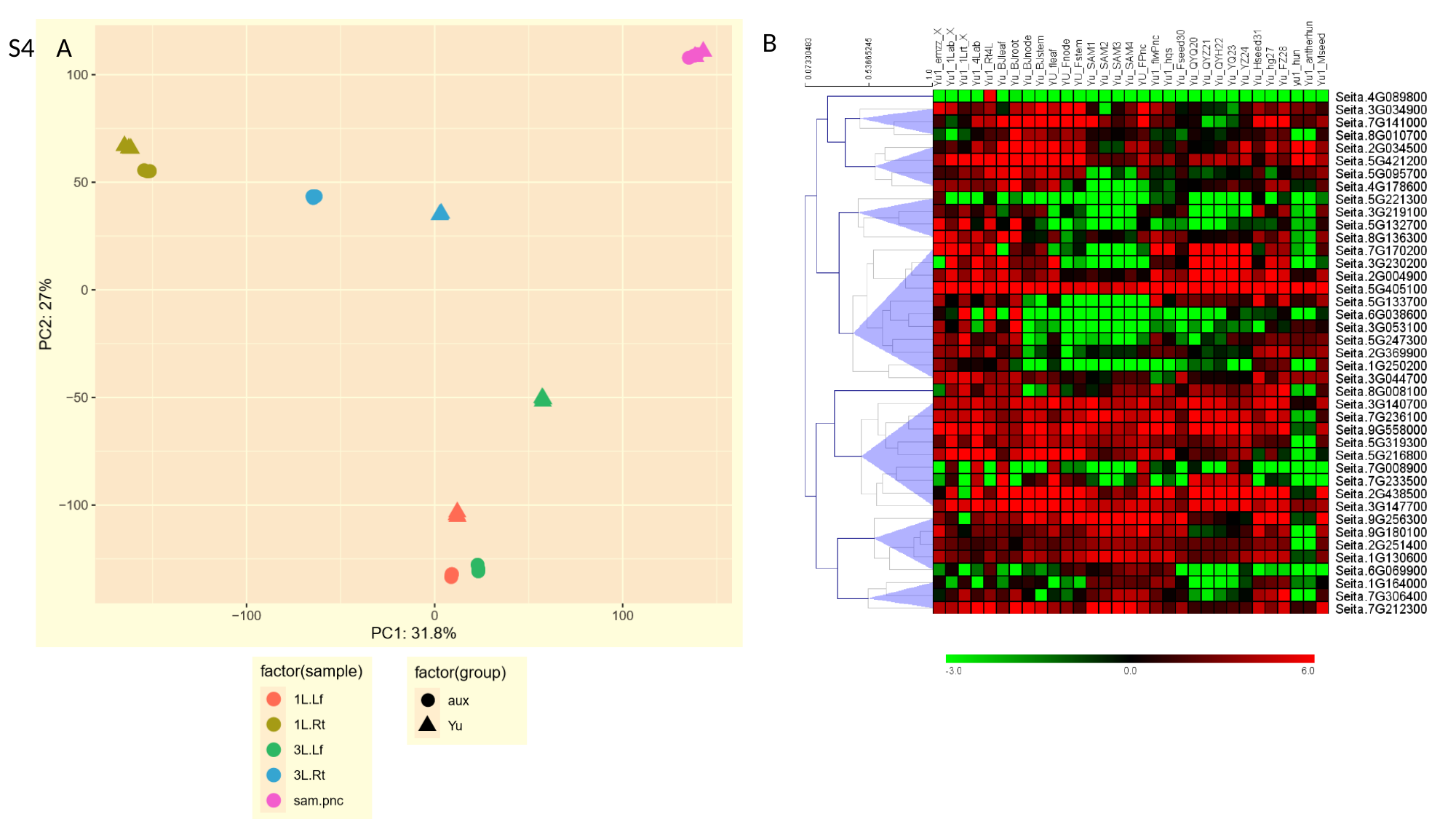

B
S4
A

### Slide 5
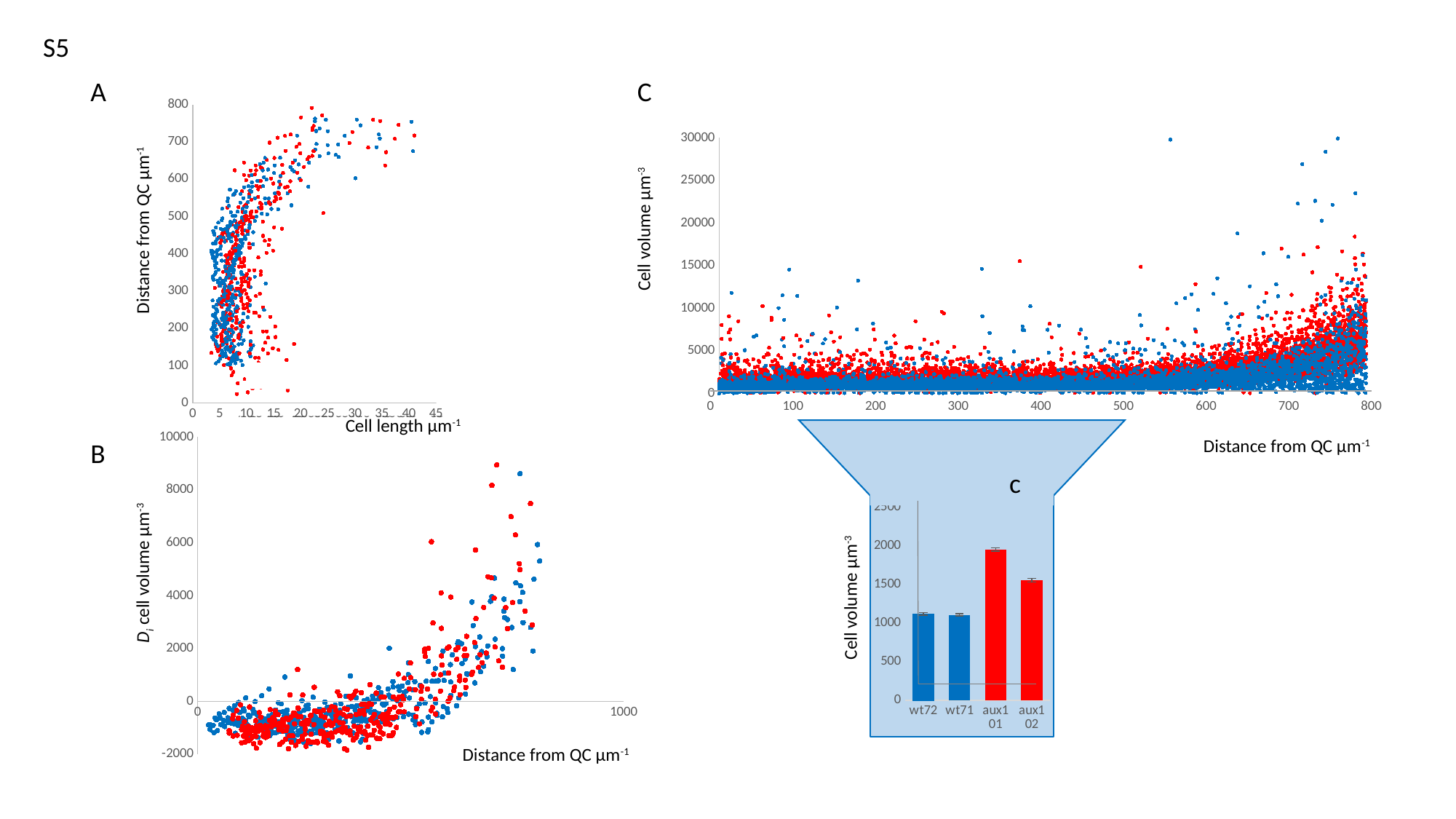

S5
C
A
#### Chart
| Category |
|---|
#### Chart
| Category |
|---|
#### Chart
| Category |
|---|
#### Chart
| Category |
|---|
#### Chart
| Category | | | | |
|---|---|---|---|---|0
100
200
300
400
500
600
700
800
Cell volume µm^-3^
Distance from QC µm^-1^
#### Chart
| Category | | | |
|---|---|---|---|Cell length µm^-1^
#### Chart
| Category | average 100 to 500 µm |
|---|---|
| wt72 | 1118.6215016891867 |
| wt71 | 1108.4964382470123 |
| aux1 01 | 1953.3702500000009 |
| aux1 02 | 1555.4741215981471 |
#### Chart
| Category |
|---|
#### Chart
| Category |
|---|
#### Chart
| Category | | | |
|---|---|---|---|Distance from QC µm^-1^
B
c
D_i_ cell volume µm^-3^
Cell volume µm^-3^
Distance from QC µm^-1^
Average cell volume in the epidermis between 100 and 500 µm from the QC
